# Combining quantitative proteomics and interactomics for a deeper insight into molecular differences between human cell lines

**DOI:** 10.1101/2024.06.12.598691

**Authors:** Anna A. Bakhtina, Helisa H. Wippel, Juan D. Chavez, James E. Bruce

## Abstract

Cellular functional pathways have evolved through selection based on fitness benefits conferred through protein intra- and inter-molecular interactions that comprise all protein conformational features and protein-protein interactions, collectively referred to as the interactome. While the interactome is regulated by proteome levels, it is also regulated independently by, post translational modification, co-factor, and ligand levels, as well as local protein environmental factors, such as osmolyte concentration, pH, ionic strength, temperature and others. In modern biomedical research, cultivatable cell lines have become an indispensable tool, with selection of optimal cell lines that exhibit specific functional profiles being critical for success in many cases. While it is clear that cell lines derived from different cell types have differential proteome levels, increased understanding of large-scale functional differences requires additional information beyond abundance level measurements, including how protein conformations and interactions are altered in certain cell types to shape functional landscapes. Here, we employed quantitative *in vivo* protein cross-linking coupled to mass spectrometry to probe large-scale protein conformational and interaction changes among three commonly employed human cell lines, HEK293, MCF-7, and HeLa cells. Isobaric quantitative Protein Interaction Reporter (iqPIR) technologies were used to obtain quantitative values of cross-linked peptides across three cell lines. These data illustrated highly reproducible (R^2^ values larger than 0.8 for all biological replicates) quantitative interactome levels across multiple biological replicates. We also measured protein abundance levels in these cells using data independent acquisition quantitative proteomics methods. Combining quantitative interactome and proteomics information allowed visualization of cell type-specific interactome changes mediated by proteome level adaptations as well as independently regulated interactome changes to gain deeper insight into possible drivers of these changes. Among the biggest detected alterations in protein interactions and conformations are changes in cytoskeletal proteins, RNA-binding proteins, chromatin remodeling complexes, mitochondrial proteins, and others. Overall, these data demonstrate the utility and reproducibility of quantitative cross-linking to study systems-level interactome variations. Moreover, these results illustrate how combined quantitative interactomics and proteomics can provide unique insight on cellular functional landscapes.

**GRAPHICAL ABSTRACT:** 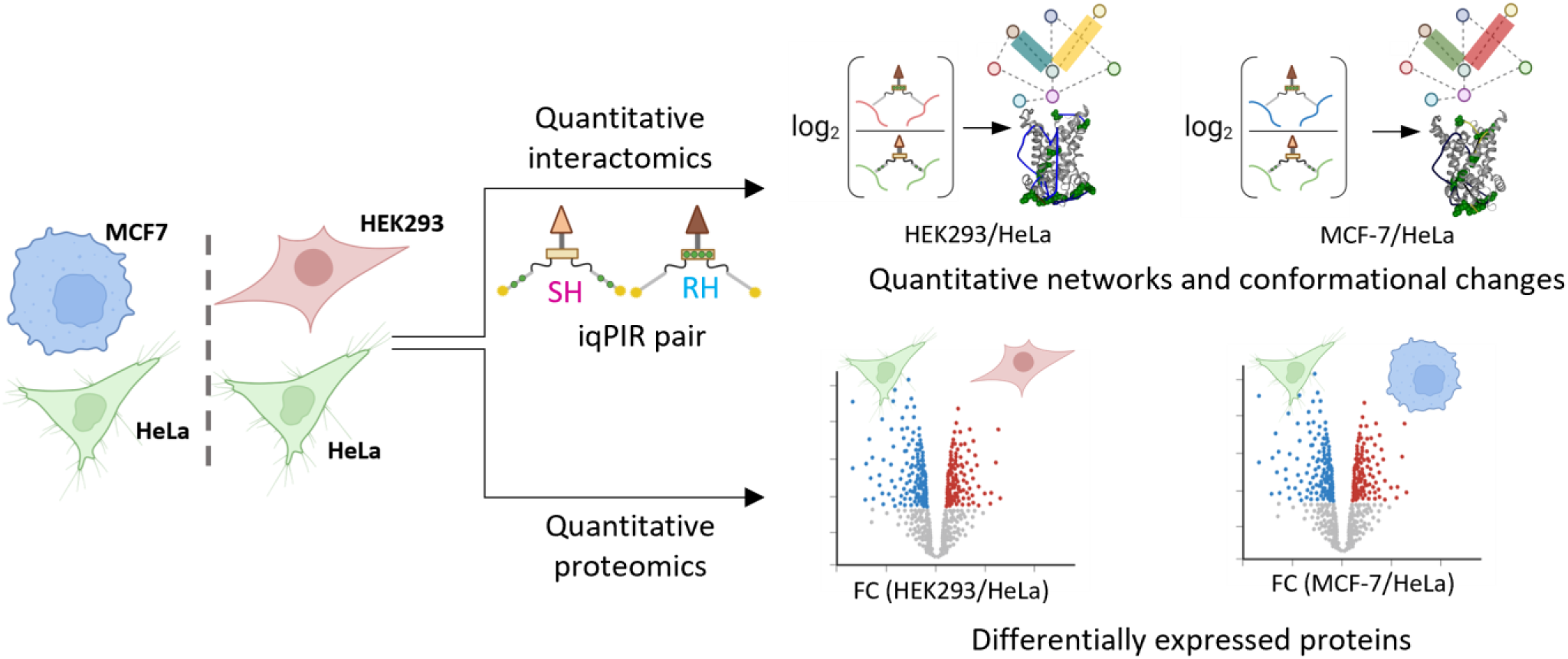

## INTRODUCTION

Human cell lines are widely used as model systems for biological research. Because these cell lines are originally derived from human tissues, the physiological and molecular alterations of these cells in response to environmental perturbations, e.g., drug treatment, are often considered a proxy to what happens in more complex living systems. Human cell lines have been used for vaccine and antibody production, drug discovery, studies on drug metabolism and cytotoxicity, and synthesis of biological compounds^1^. Furthermore, the information provided by biological studies using cell lines as a model is key to understanding molecular mechanisms of human diseases. Among the most commonly used human cell lines are the cervical cancer cell line HeLa, breast cancer MCF-7, and immortalized embryonic kidney HEK293. Albeit originating from the same species, these cell lines present cell-specific differences on the molecular level, including varying protein abundance levels^2^, different protein-protein interactions (PPIs)^3^, conformations, and function. These cellular alterations need to be considered when selecting the best model for a study. To date, some studies have applied mass spectrometry (MS)-based proteomics to compare the protein content of human cell lines and pinpoint differences in the proteomic profiles among them^2,4,5^. Protein abundance levels constitute only one of many regulators of the functional landscape that exists within cells. Large scale profiling by affinity-purification MS (AP-MS) has revealed extensive customization of the PPIs across different cell lines^3^. Large-scale AP-MS requires genetic alteration of cells to exogenously express affinity tagged bait proteins of interest, which can alter the native cellular environment and interactions of some proteins. Moreover, little or no structural information is gained on protein interactions and conformations from most AP-MS studies. On the other hand, chemical cross-linking of proteins coupled to mass spectrometry (XL-MS) is an emerging tool to study protein structures and the interactome^6–8^. XL-MS is based on the reactivity of cross-linker groups toward specific side chains of proteins, usually primary amines. The distance between these reactive groups serves as a physical constraint that requires cross-linked sites exist proximal to one another and is useful for modeling protein structures and protein complex topologies^8^. While the structural information gained by XL-MS is lower resolution than traditional structural biology techniques such as X-ray crystallography and cryo-EM, the information derived from XL-MS is highly complementary^7,9^ and can be combined with AlphaFold3 to advance large-scale protein-protein structure predictions^10,11^. Furthermore, XL-MS is applicable to intact living cells and tissues devoid of any genetic alteration, thereby providing structural insight on proteins and complexes in their native environments.

When applied quantitatively, qXL-MS^12^ offers unique insight on protein conformational and protein-protein interaction changes inside cells^13^ and even in animal phenotypic comparisons^14,15^. The direct quantitation of each specific pair of lysine residues cross-linked under different conditions allows large-scale visualization of changes in molecular interactions within complex systems not possible by any other means. Combining qXL-MS data with traditional quantitative proteomics methods to obtain information about changes in protein abundance levels promotes deeper understanding of the regulators that drive functional changes inside living systems. Here, we applied *in vivo* qXL-MS using quantitative isobaric Protein Interaction Reporter (iqPIR)^16^ technologies to perform a comparison of three commonly used human cell lines: HEK293, MCF-7, and HeLa. This study enabled pairwise comparison of HEK293 and MCF-7 with HeLa. The iqPIR strategy allows mixing equal total protein amounts from compared samples before proteolytic digestion, decreasing variability due to subsequent sample processing. These efforts revealed the first large-scale changes in protein conformations and interactions that relate to functional changes among the three commonly studied human cell lines. The qXL-MS results also demonstrate that quantitation values from iqPIR are highly reproducible with all biological replicates and changes as low as 0.5 on a log_2_ scale can be detected with statistical significance. Highlighted proteins with changing cross-link ratios in different cell lines include keratins, RNA-binding proteins, chromatin remodelers, proteins involved in clearance of cytotoxic metabolites, and mitochondrial proteins. Leveraging quantitative proteomics enables the proposal of possible drivers of the change in the cross-link levels.

## RESULTS and DISCUSSION

Quantitative cross-linking experiments resulted in the identification of 1,797 cross-linked peptides across two biological replicate comparisons of MCF-7 and HeLa (MCF-7/HeLa), and 2,361 cross-links across 4 biological replicate comparisons of HEK293 and HeLa (HEK293/HeLa). The complete cross-linking information is made available at Supplementary Table 1 and online on XLinkDB at http://xlinkdb.gs.washington.edu/xlinkdb/PHP_SAVED_VIEWS/anna/HEK293_HeLa_and_MCF7_HeLa_iqPIR_cell_lines_interactomics.php. After imposing quality filters for each quantitative value (95% confidence ≤ 0.5 and at least 4 ions used for calculation of log_2_ ratio) there are 982, 851, 1,020, 1,158, and 878, 853 high confidence quantitative values in each biological replicate of HEK293/HeLa and MCF-7/HeLa, respectively (**Figure 1A**). The distribution of cross-linked peptide log_2_ ratios appears normal, with the large portion of cross-link levels not changing between the cell lines (**Figure 1B**). It should be emphasized that all interactome quantitative values are derived from cross-link levels produced inside live cells prior to any protein extraction. Unlike quantitative proteome measurements that derive abundance information based on extracted protein levels, any differences that may or may not occur in protein extraction from one cell sample to the next in iqPIR experiments are accounted for by mixing iqPIR-labeled samples with 1:1 total protein amounts and further by centering the iqPIR log_2_ ratio distribution to zero if needed (**Supp. Figure 1A**). Thus, iqPIR quantitative interactome data are unique in that the derived ratios are reflective of changes that exist inside cells in terms of the amount of each cross-link level produced prior to lysis and protein extraction. As such, iqPIR ratios are affected by protein modification, conformational, and interaction levels that exist within the cellular environment and show cell type-specific changes as illustrated for the first time below. Moreover, the iqPIR method shows great reproducibility across multiple biological replicates, perhaps due to tight regulatory control on the intra-cellular environment where cross-linking reactions are performed. In both HEK293/HeLa and MCF-7/HeLa datasets, high correlation among forward and reverse labeling biological replicate samples was observed, with Pearson’s correlation coefficient R^2^ values of 0.91 and 0.88 respectively (**Figure 1 C,D**). On the other hand, comparison of quantitative values between a forward replicate of HEK293/HeLa and a forward replicate of MCF-7/HeLa showed much weaker correlation, as expected (R^2^ = 0.01; **Supp. Figure 1B**). To obtain a single log_2_ ratio for each cross-link species in all HEK293/Hela or MCF-7/HeLa data, all ions that provided quantitative information (i.e., released peptides and fragment ions containing cross-linked lysine sites which provided iqPIR isotopic signatures) coming from all biological and technical replicates were combined. Although differential amino acid modifications can affect protein conformations, trypsin cleavage, and intra-protein cross-link levels, most cross-linked species involving linkages of the same lysine residues but differing peptide pairs, such as those arising due to trypsin missed cleavage sites and/or methionine oxidation, are anticipated to provide quantitation consistent with fully cleaved, unmodified products that have redundant linkage. The data acquired in these experiments indeed illustrate this to be the case as shown in **Supp. Figure 1C** and provide further increased confidence in iqPIR derived quantitation of cross-linked levels. Allowing a maximum of one missing quantitative value out of four biological replicates resulted in 629 high confidence quantified cross-linked species in the HEK293 to HeLa interactome dataset. Filtering of MCF-7/HeLa interactome data for zero missing values among both biological replicates resulted in 715 high confidence quantified cross-linked species. Application of Student’s *t*-test with Bonferroni multiple testing correction and log_2_ ratio greater than 0.5 or less than -0.5 resulted in 285 HEK293 cross-links and 382 MCF-7 cross-links with log_2_ ratios significantly different from HeLa (**Figure 1E,F**; **Supp. Table 2**). Among all cross-linked peptide pairs, 545 are common among all comparisons (**Supp. Figure 1D**). We then compared protein levels among the cell lines with DIA-Based quantitative proteomics. As expected, proteomes of HEK293, MCF-7, and HeLa are quite distinct from each other with many protein levels significantly different between them (**Supp. Figure 1E-G**). Comparison of intra- link levels, where cross-linked peptides originate from the same protein, with protein abundance changes from the proteomics data can reveal cross-linked peptide level changes that are driven by proteome-level changes, and those that are independent of protein-level change. Comparison of high confidence intra-link and relevant protein level changes revealed correlation between intra-link and protein levels (HEK293/HeLa R^2^ = 0.48 and MCF-7/HeLa R^2^ = 0.66, **Figure 1G**), albeit weaker correlation than observed between replicates of either interactome datasets (**Figure 2C,D**). This result is expected since changes in protein levels are likely to result in changes in cross-link levels. Cross-link level changes that differ from protein level changes indicate differences in lysine accessibility, reactivity, or site-to-site relative proximity, such as those due to interaction, modification or conformational differences that may be present in different cell types.

**Figure 1.**
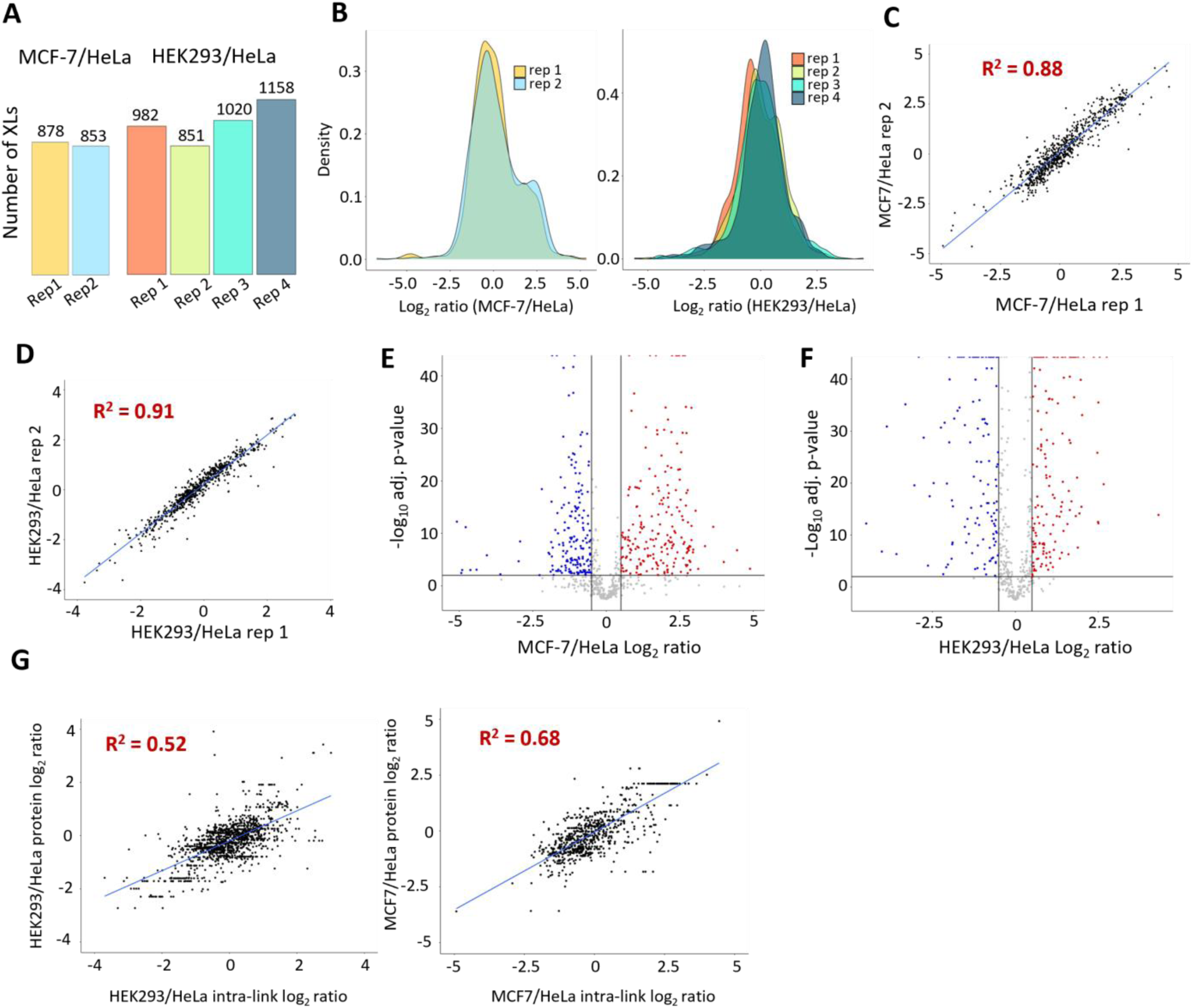
Quantitative interactomics with iqPIR. **A**. High confidence cross-links quantified in each biological replicate. **B**. Distribution of log_2_ ratios of quantified cross-links in each biological replicate. **C**. Correlation between biological replicates of MCF-7 and HeLa comparison, Pearson’s R^2^ = 0.88. **D**. Correlation between biological replicates of HEK293 and HeLa comparison, Pearson’s R^2^ = 0.91. Volcano plots with log_2_ ratio of cross-links quantified with 95% confidence ≤ 0.5 in both biological replicates in MCF-7 to HeLa comparison (**E**) or with 95% confidence ≤ 0.5 in at least 3 distinct biological replicates in HEK293 and HeLa comparison (**F**) based on all contributing ions. Bonferroni corrected *p*-value of 0.05 and |log_2_ FC| > 0.5 are used to indicate significance. **G.** Correlation between log_2_ ratios for intra-protein cross-links and respective log_2_ ratios for proteins based on whole proteome quantitation for HEK293 and HeLa comparison with R^2^ = 0.52 and MCF-7 and HeLa comparison with R^2^ = 0.68.

**Figure 2.**
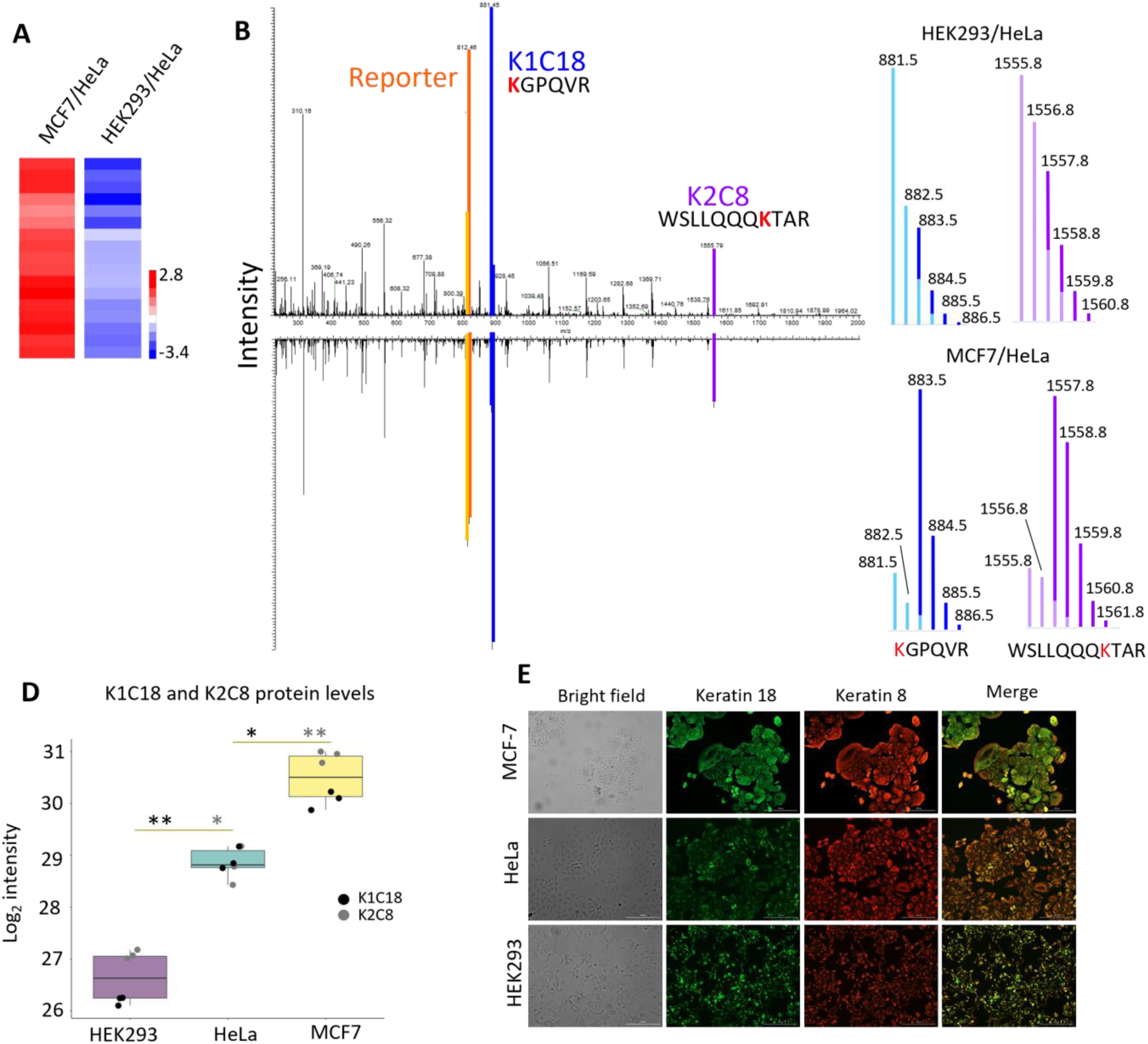
Keratins and their interactions are differentially regulated in HEK293 and MCF-7 compared to HeLa cells. **A**. Heatmap of log_2_ ratios of keratin cross-link levels produced in live cells showing that all K2C8 and K1C18 intra- and inter-link levels in HEK293 and MCF-7 cells show opposing changes, relative to link levels in HeLa cells. **B**. Spectra of a cross-link between K119 on keratin 18 and K130 on keratin 8 in HEK293/HeLa and MCF-7/HeLa are displayed as mirror images. Peaks for reporter ions (orange), peptide A (blue), and peptide B (purple) are shown on the original spectra. **C**. Isotopic envelops with the apportionment from RH and stump SH for peptides A and B show the opposite direction of change in HEK293 cells and MCF-7 cells compared to HeLa. In HEK293 (cross-linked with SH), the cross-link level is decreased relative to HeLa (cross-linked with RH). In MCF-7 (cross-linked with SH), the same cross-link shows the opposite change relative to HeLa (cross-linked with RH). In all plots, darker shaded color indicates apportioned intensity from SH sample, and lighter shading indicates RH samples. **D**. Boxplot of log_2_ intensities for keratin 18 and keratin 8 protein levels in all three cell lines with significance as determined by Student’s *t*-test with multiple testing correction (* = 0.05, ** = 0.01). Each point indicates a value for either K1C18 (black) or K2C8 (grey) in each biological replicate. **E**. Immunofluorescence imaging with anti-keratin 18 (green) and anti-keratin 8 (red) antibodies in all cell lines indicating protein level differences consistent with proteome measurements.

### Interactome changes driven by proteome-level changes

A key advantage of combining quantitative interactome and proteome datasets is the opportunity to further evaluate novel quantitative interactome technologies against more widely established quantitative methods in cases where the changes should be coordinated. Through pathways analysis of proteins with differential cross-links, we identified cytoskeleton organization as one of the pathways significantly different in both datasets (**Supp. Table 3**). Keratin proteins belong to this pathway and have one of the biggest changes in both datasets. Keratins are intermediate filament proteins that form heterodimers of type 1 (keratins 9-28) and type 2 (keratins 1-8 and 71-80) and can be used as a prognostic tool in many types of cancer^17^. Type 2 keratin 8 (K2C8) and type 1 keratin 18 (K1C18) exist as heterodimers and often exhibit protein abundance interdependent upon one another ^17,18^. Thus, proteome changes of either protein are expected to be coordinated with level changes of the partner as well as changes in the K2C8-K1C18 interaction levels. In MCF-7/HeLa K2C8 and K1C18 intra-link and inter-link levels are significantly increased (mean log_2_ ratio of 2.5), while in HEK293/HeLa interactome data these same link levels are significantly decreased by nearly the same amount (mean log_2_ ratio of -2.9) (**Figure 2A**). Previous work in our lab showed that keratin cross-link levels change during the paclitaxel treatment of HeLa cells and are also a hallmark of chemoresistance^19,20^. With paclitaxel treatment^20^ as well as in chemoresistant carcinoma cells^19^ the cross-link between K1C18 lysine 119 (K119) and K2C8 K130 was observed to be increased in both cases. With iqPIR technologies, multiple ions from both cross-linked peptides provide quantitative information relevant to this interaction. For example, with the K1C18 K119-K130 K2C8 cross-link, we have 66 and 37 measurements of ratios for peptide and fragment ions in MCF-7/HeLa and HEK293/HeLa, respectively (**Supp. Figure 2A,B**). Representative spectra from each interactome analysis, both coming from the experiments where HeLa is cross-linked with reporter heavy (RH) iqPIR cross-linker, so HeLa peptides carry a 2 Da lighter stump than either HEK293 or MCF-7, show remarkable similarity, despite opposite ratio measurements (**Figure 2B**). Interactome quantitation is done on the MS^2^ level after the cross-linker is cleaved from the peptides by decomposing peaks in isotopic envelopes of peptides and their fragments into contributions from RH and stump heavy (SH) cross-linked samples (**Figure 2C, Supp. Figure 2C-F**). The iqPIR reporter can in principle also be used for iqPIR quantitation, but because it is the same for all cross-linked species, it can suffer from ratio compression similar to other isobaric quantitative methods^21^. For this reason, iqPIR reporter ion intensities are not included when calculating the cross-linked peptide ratios (**Supp. Figure 2G**). Quantitative proteome-level analyses revealed that K2C8 and K1C18 protein levels are lower in HEK293 and higher in MCF-7 compared to HeLa (**Figure 2D**), log_2_ fold change of -1.6 (adj. p-value = 0.033) and -2.6 (adj. p-value = 0.002) in HEK293/HeLa and 2.15 (adj. p-value = 0.0095) and 1.18 (adj. p-value = 0.017) in MCF-7/HeLa. These data indicate generally good agreement between interactome and proteome quantitation expected for K2C8-K1C18 and further validate iqPIR quantitative capabilities. We also performed fluorescence imaging of MCF-7, HEK293, and HeLa using anti-keratin antibodies (**Figure 2E**). While not directly quantitative, the images also show the same trend of keratins being more abundant in MCF-7 than in HEK293 cells. Thus, iqPIR data yield expected changes in interactome levels based on measured proteome and immunofluorescence data.

### Quantitative interactomics allows a deeper view of remodeled protein networks

One of the key advantages of interactome analysis is the potential to reveal molecular details and alterations that are not driven by protein abundance level changes that may be difficult to observe otherwise. Pathway analysis of proteins with differential cross-links showed that the most significant molecular function for both datasets was RNA binding (**Supp. Table 3** and **Supp. Figure 3**). RNA-binding proteins (RBPs) interact with RNAs to regulate many key cellular processes, e.g., transcription, splicing, mRNA localization, transport, and translation^22^. Heterogeneous nuclear ribonucleoproteins (hnRNPs) are RBPs that contain their own nuclear localization signal and therefore comprise complexes with RNA primarily within the nucleus^23–25^ and hnRNPs exist under complex functional regulatory control based on cellular demands. Interactome quantitation revealed large changes in cross-link levels of many hnRNPs in both HEK293/HeLa and MCF-7/HeLa datasets. A total of 61 and 102 cross-links were detected for hnRNPs in MCF-7/HeLa and HEK293/HeLa interactome datasets, respectively (**Supp. Table 4**). A quantitative network that encompasses cross-links from 19 RNA binding proteins was constructed allowing visualization of cross-linked peptide levels that change in both MCF-7 and HEK293, only one cell comparison, or did not change (**Supp. Figure 4A**). Proteome-level analysis of these proteins indicates that some exhibit levels that are changed in HEK293 and/or MCF-7 relative to HeLa cells and thus, are expected to exhibit altered cross-link levels (**Supp. Figure 4 B,C**). Overall, distributions of protein level log_2_ fold changes show that MCF-7/HeLa values are shifted toward negative and HEK293/HeLa are shifted toward positives compared to distributions of all XLs or proteins (**Figure 3A, Supp. Figure 4D**). A similar trend is observed on the cross-link level, although with a more pronounced shift, suggesting that even greater remodeling happens on the interactome level.

**Figure 3.**
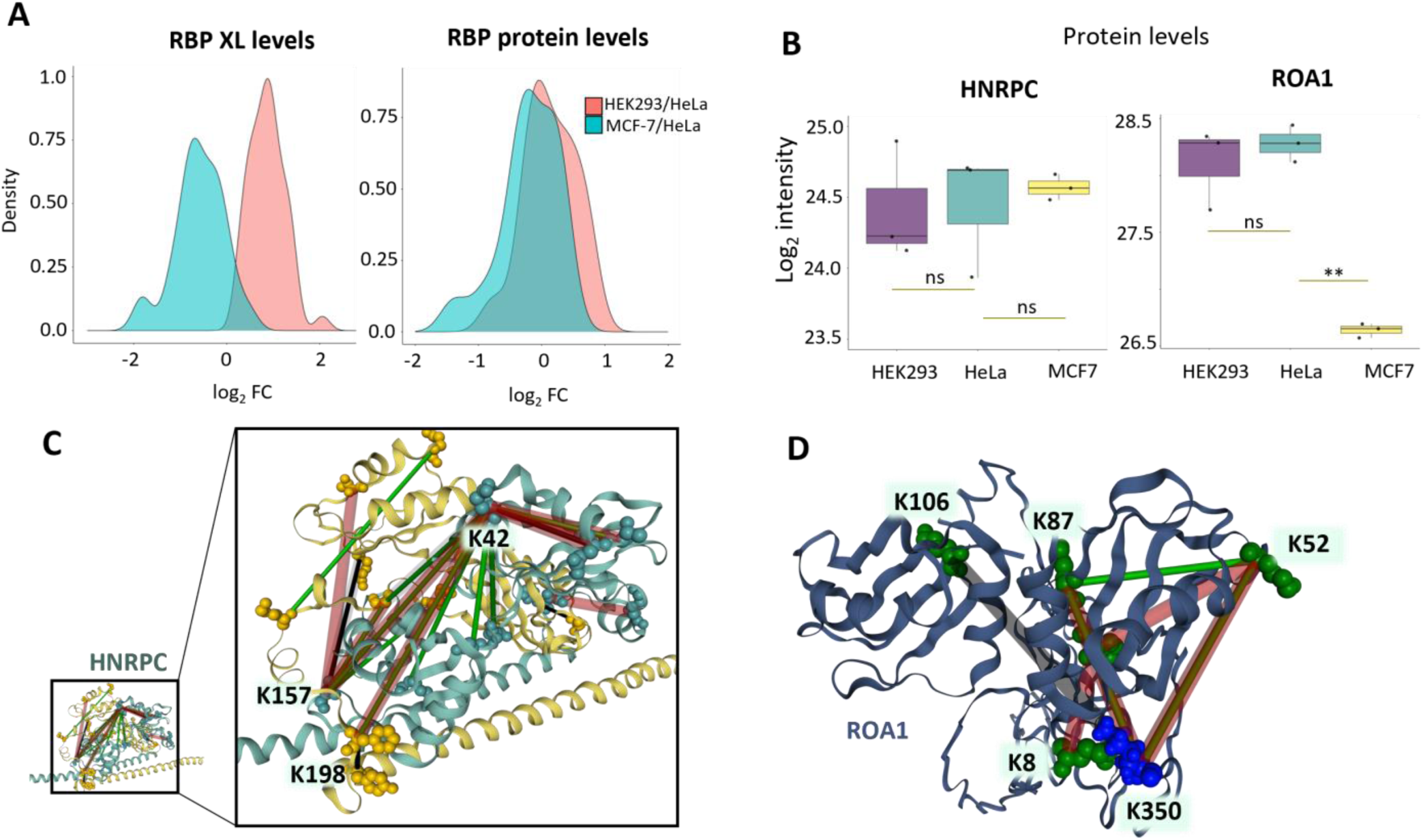
Proteome and interactome level remodeling of RNA binding proteins. **A.** Distributions of average log_2_ ratios of RNA binding proteins on a cross-link level (left) and protein level (right). **B.** Boxplots of log_2_ intensities of HNRPC and ROA1 protein levels in all three cell lines with significance, as determined by Student’s *t*-test with multiple testing correction (** = 0.01, ns = non-significant). **C**. Cross-links for HNRNPC, mapped to the SFPQ crystal structure (PDB 4WIJ). Cross-links from HEK293/HeLa and MCF-7/HeLa are indicated with thick transparent lines and thin solid lines, respectively. The log_2_ ratio intensities are represented in green for decreasing ratios compared to control, in red for increasing ratios, or black for non-changing ratios. **D.** Cross-linked peptides for ROA1 mapped to a previously described modeled structure^29^. Lysine K350, indicated in blue, is part of the nuclear targeting sequence M9.

Among the proteins exhibiting cross-link level changes that differ from protein level changes are HNRPC (similar protein levels in all three cell lines) and ROA1, which exhibits similar protein levels in HEK293 and HeLa, but is repressed in MCF-7 cells (**Figure 3B**). HNRPC is involved in binding pre-mRNA and has a role in splicing by binding regulatory untranslated regions^26^. ROA1 is involved in RNA packaging and has been found to be antagonistic toward exon usage in splicing^27^. Previous mass spectrometry studies identified peptides indicating both HNRPC and ROA1 may be components in isolated nuclear spliceosomal B and C complexes^28^, although neither protein has yet been structurally characterized within resolved spliceosome complexes. The majority of HNRPC intra-links are increased in HEK293 and decreased in MCF-7, when compared to HeLa cells (**Figure 3C**). The links are mapped to an SFPQ structure as previously described as there is no solved HNRPC structure^29^. Other HNRPC intra-link levels were observed with levels that were unaltered in either interactome dataset, which is expected for and consistent with unaltered HNRPC protein levels (**Supp. Table 4**). Combining these results with observed unaltered protein levels indicates that the altered HNRPC cross-link levels reflect molecular changes due to site accessibility, reactivity, or relative site-site proximity inside these cells not driven by differential protein abundance levels. These observations must therefore arise due to differential changes in modifications, conformations, and/or molecular interactions of HNRPC within these three cell lines.

Moreover, inter-protein cross-links between HNRPC and ROA1 were also observed with altered levels in these inter-protein comparisons (Supp. Figure 4A). While all HNRPC-ROA1 inter-link levels decreased in MCF-7/HeLa data (log_2_ ratios of -1.79 for K42-K350 and -1.84 for K157-K350), these same links were observed with increase levels in the HEK293/HeLa interactome with log_2_ ratios of 0.3 for K42-K350, and 1.1 for K157-K350, with the latter passing applied significance and ratio filters. Protein levels of ROA1 decreased in the MCF-7/HeLa proteome and thus, likely contribute to and may explain the decreased levels of ROA1 intra- and inter-link levels (**Figure 3D**). However, neither HNRPC nor ROA1 protein levels changed significantly in the HEK293/HeLa proteome comparison. Therefore, other molecular differences as discussed above must give rise to increased HNRPC-ROA1 inter-linked levels in HEK293/HeLa cell interactome changes. Interestingly, all HNRPC-ROA1 inter-linked peptides and all inter- links between ROA1 and all other hnRNPs other than ROA2 involve the single ROA1 lysine K350 (**Supp. Figure 3A**). This residue falls within the nuclear targeting sequence M9 (residues 320-357), involved in the transport and localization of mRNAs^30^. The role of K350 in hnRNPs conformation and protein-protein interactions has been previously observed^29^. The present quantitative interactome data indicate that ROA1 M9 involvement in mRNA nuclear shuttling is different in HEK293 compared to HeLa cells in ways that are not regulated through ROA1 protein levels.

### Chromatin remodeling complexes and their interaction with nucleosomes is altered between HEK293 and HeLa

Epigenetic landscape and chromatin remodeling has been shown to play an important role in cancer onset and progression^31^. There are several important families of chromatin remodelers that have been characterized extensively. One of them is ISWI family, first discovered in drosophila^32^. A characteristic feature of a catalytic subunit of ISWI remodelers is SANT domain^33^. This domain, together with a neighboring SLIDE domain, is responsible for nucleosome recognition. We have identified and quantified a cross-link (K847-K855) that spans SA1 helix of the SANT domain of SMCA5, a human ISWI catalytic subunit encoded by the SMARCA5 gene (**Figure 4A, Supp Figure 5A**). In HEK293/HeLa, this cross-link is increased, while it is unchanged in MCF-7/HeLa. There are also no protein level differences between any of the cell lines, so it is unlikely that changes in HEK293/HeLa are caused by protein abundances (**Figure 4B**). We have mapped this cross-link on a structure of the SANT domain of *D. Melanogaster* homologue of SMCA5, ISWI (**Figure 4C left**), but there is no solved structure of SMCA5 that contains the SANT domain. Recently, a structure of *S. Cerevisiae* ISW1a complexed with two nucleosomes have been solved^34^. The sequence of yeast ISW1a is substantially divergent from fly ISWI (and human SMCA5), but structurally their SANT domains are similar (**Supp. Figure 5B**). Mapping cross-linked residues on this structure shows that the K847-K855 cross-link would be right at the contact with DNA connecting two nucleosomes (**Figure 4C right**). Together with non-changing protein levels we can hypothesize that increase in the cross-link is caused by the decreased interaction with DNA and nucleosomes in HEK293 cell line. Chromatin accessibility and transcriptional machinery differences between the human cell lines have been reported before^35^ and present work illustrates how quantitative interactomics can help in their investigation.

**Figure 4.**
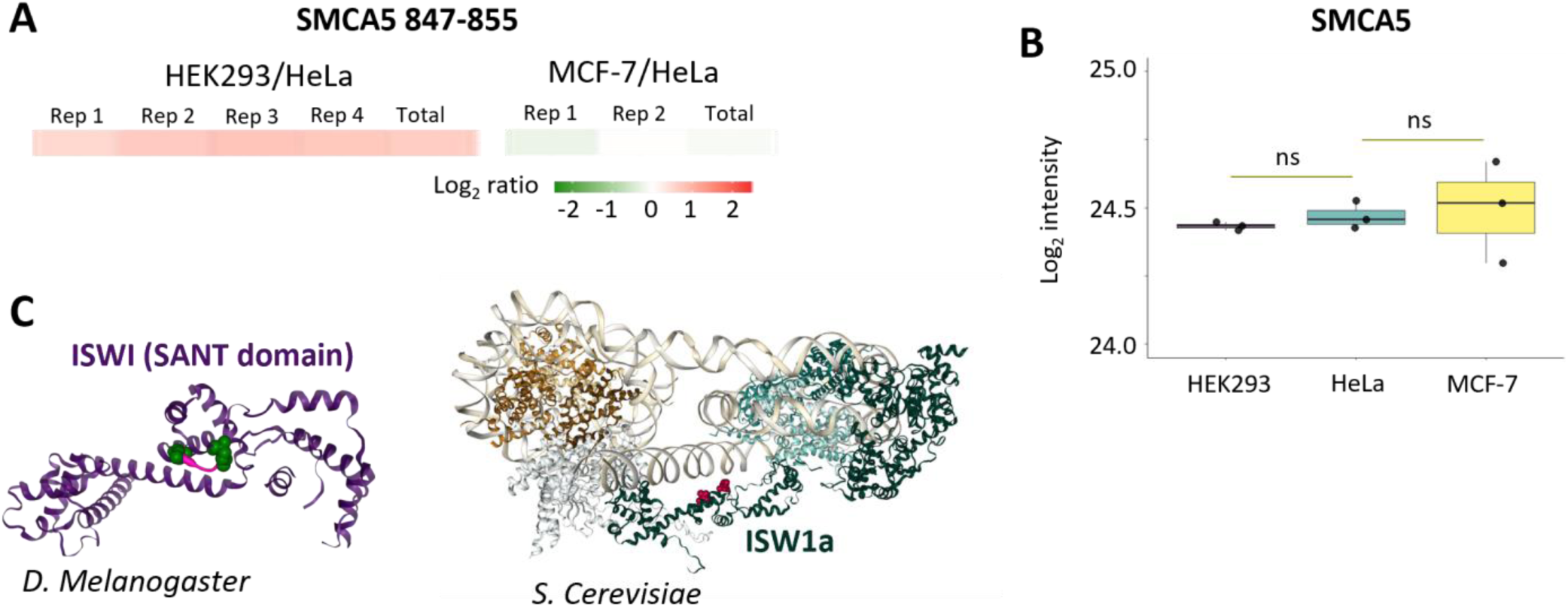
Chromatin remodeling complexes interaction with nucleosome is different in HEK293 compared to HeLa. **A.** Heatmap of log_2_ ratio of SMCA5 K847-K855 intra-link. **B**. Boxplot of SMCA5 log_2_ intensities in HEK293, MCF-7 and HeLa with significance as determined by Student’s *t*-test with multiple testing correction (ns = non-significant). **C**. SMCA5 cross-link mapped to the structure of fly SANT domain (PDB:1OFC) (left) and cross-linked residues (magenta) mapped on the structure of yeast ISWE1a complexed with dinucleosomes (PDB:7X3T) (right).

**Figure 5.**
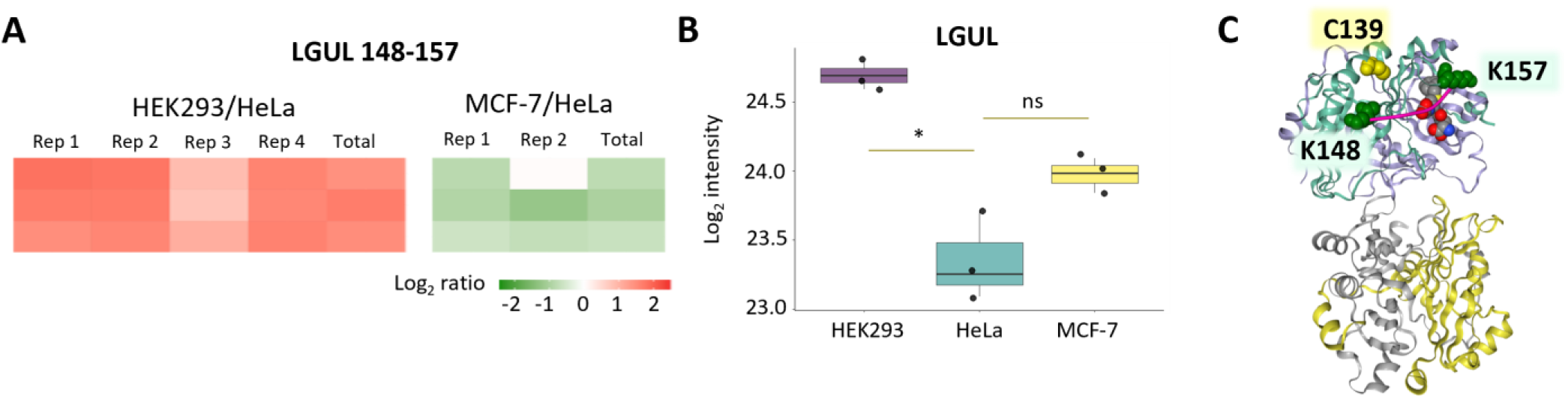
Glyoxolase 1 (LGUL) cross-link at the regulatory site is decreased in MCF-7/HeLa. **A.** Heatmap of log_2_ ratios of LGUL K148-K157 cross-links in HEK293/HeLa and MCF-7/HeLa. **B.** Boxplot of LGUL log_2_ protein intensities in HEK293, MCF-7, and HeLa with significance as determined by Student’s *t*-test with multiple testing correction (* = 0.05, ns = non-significant). **C.** LGUL K148-K157 cross-link mapped to a human structure with an S-benzyl-glutathione inhibitor (located between cross-linked residues) (1FRO). Cysteine modified by inhibitory glutathionylation is indicated in yellow.

### Interactome changes in an enzyme that protects against cytotoxic glycolytic byproducts

Protein glyoxalase 1 (LGUL) is encoded by GLO1 gene and is a component of the system responsible for detoxification of cytotoxic compound methylglyoxal (MG) by the reaction with glutathione (GSH)^36^. MG is a highly reactive byproduct of glycolysis and a precursor of glycation end-products, associated with multiple pathologies^37^. The switch to a glycolytic phenotype is one of the main features of most cancers that results in accumulation of cytotoxic MG ^38,39^. Overall, overexpression or increased activity of LGUL has been reported in many cancers and is associated with poor prognosis^39,40^. Quantification of the LGUL K147-K158 cross-link level was achieved with multiple peptides resulting from missed cleavage and methionine oxidation in both HEK293/HeLa and MCF-7/HeLa datasets (**Figure 5A**). Increase in this cross-link in HEK293 cells appears to be driven by increased protein levels (**Figure 5B**). Unlike HEK293/HeLa, in MCF-7/HeLa this cross-link changes in the opposite direction: cross-link level is decreased while protein levels slightly increase (no statistical significance is reached). Mapping this cross-link on the structure with a GSH analogue that acts as competitive inhibitor shows that the cross-link spans the GSH binding pocket (**Figure 5C**). Moreover, this site is proximal to cysteine 139 (C139), which has been reported as a site of S-glutathionylation that has an inhibitory effect on LGUL activity^41^. Both C139 modification and GSH binding could potentially affect the formation of the cross-link. Molecular dynamic simulations have shown that C139 glutathionylation might affect the flexible loop where K157, involved in the cross-link discussed here, is located ^41^. Overall, these data suggest LGUL is differentially regulated in HeLa and MCF-7 cells in ways not driven by protein levels.

### Integration of quantitative cross-linking with proteomic analysis reveals dynamic functional states of adenine nucleotide translocase isoforms

Adenine nucleotide translocase (ADT) is a dual gated inner mitochondrial membrane transporter for moving ADP into the matrix and ATP into the intermembrane space in mitochondria^42^. Cross-linked peptide levels from 2 out of 4 human isoforms, ADT2 and ADT3 were quantified in these experiments. In MCF-7/HeLa we observe increase in ADT2/3 cross-link levels between K147 and K272 or K33 (**Figure 6A**). As for protein levels, there is no change in the levels of ADT2 and a decrease in ADT3 in MCF-7 compared to HeLa (**Figure 6B**). To further investigate the functional change in ADT proteins that could be associated with these cross-links we performed iqPIR experiments on the isolated mitochondria treated with two different ADT inhibitors. The mouse cell line c2c12 was employed for these studies due to the high mitochondrial content in these cells and ANT expression and high homology between human and mouse ADT proteins (**Supp. Figure 5C**). Bongkrekic acid (BKA) locks ADT in its matrix open state or m-state while carboxyatractyloside (CATR) locks it in the c-state^42,43^ open to the intermembrane space. The K147-K33 link when mapped on a c-state structure shows a solvent assessable surface distance (SASD) more than 99 Å, indicating that this link is not likely to be formed while ADT is in c-state, while it can be formed in m-state (**Figure 6C**). In agreement with this prediction, the K147-K33 link increases with BKA treatment and decreases with CATR (**Figure 6D**). SASD for the K147-K272 link when it is mapped on either BKA locked ADT structure (6GCI) or CATR structure (2C3E) shows plausible distances (38 and 34 Å, respectively), so this link could be associated with either m-state or c-state. However, the K147-K272 link levels follow the same patterns with BKA and CATR treatment of isolated mitochondria as the K147-K33 cross-link, indicating this link is increased in the m-state. Interestingly, for ADT3, 147-272 cross-link shows a very small increase. But when decreased protein levels of ADT3 in MCF-7 are taken into account, K147-K272 levels in ADT3 actually increase relative to protein levels, indicating increased enrichment in ADT3 m-state similar to that observed with ADT2. In HEK293, protein levels for both ADT2 and ADT3 are decreased relative to HeLa (**Figure 6A**). The only cross-links that are different from protein levels are the K147-K63 link in both ADT2 and ADT3 and the K147-K147 ADT2 dimer link. K147-K63 increases in both BKA and CATR treated mitochondria. Therefore, the K147-K63 link is unlikely to be associated with a particular state but might be an indicator of general lower activity of ADT. The importance of ADT and its conformational states in relationship to tumorigenesis and chemotherapy resistance has been explored before^44,45^. In the future, quantitative interactomics and proteomics can help elucidate ADT mechanistic and the metabolic role of ADT in cancer pathology.

**Figure 6.**
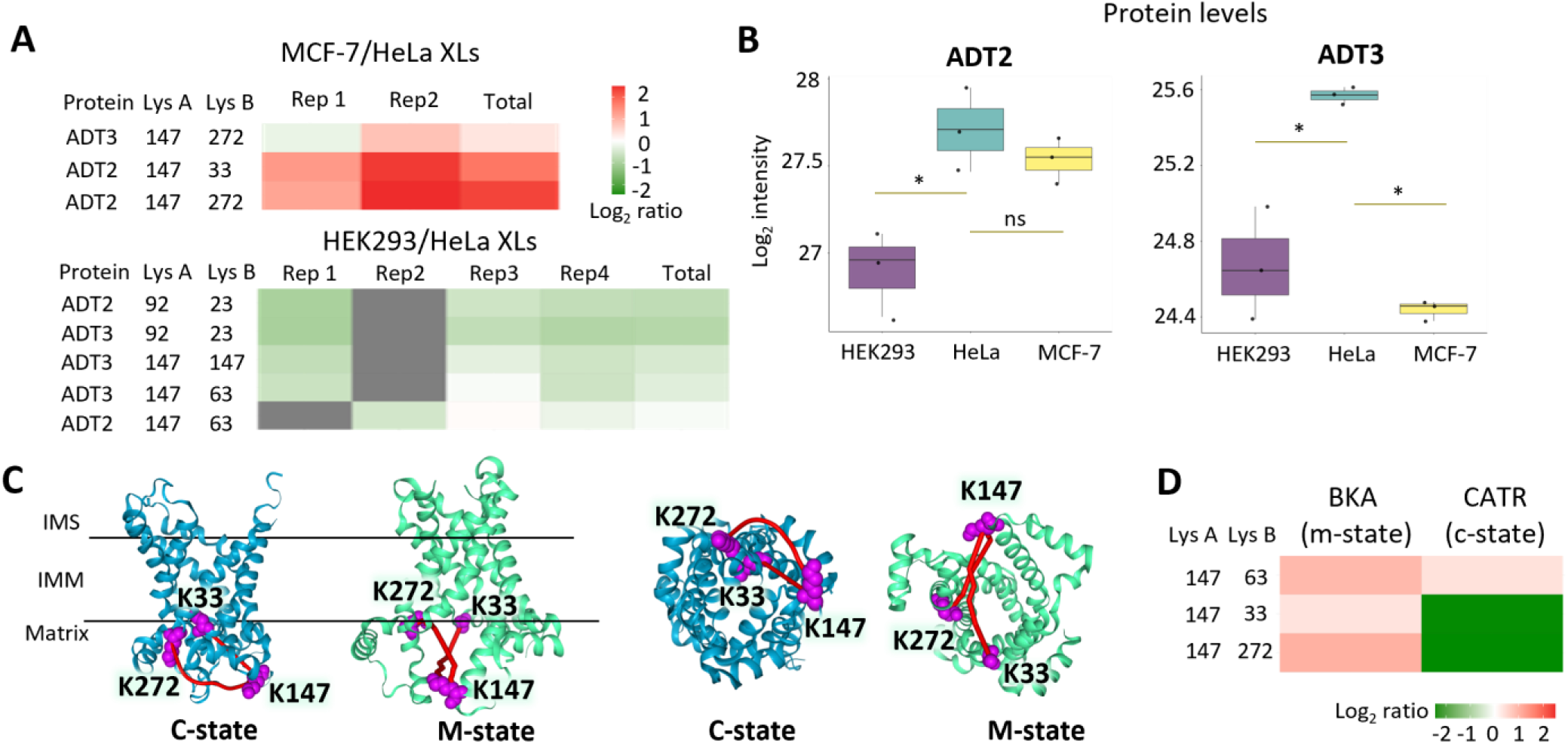
Combining quantitative interactomics and proteomics gives deeper insights into molecular differences in human cell lines. **A**. Heatmap of log_2_ ratios of ADT2 and ADT3 cross-links for MCF-7/HeLa (top) and HEK293/HeLa (bottom). **B.** Boxplot of ADT2 and ADT3 log_2_ intensities in HEK293, MCF-7, and HeLa with significance as determined by Student’s t-test with multiple testing correction (* = 0.05, ns = non-significant). Right: View from the mitochondrial matrix side. **C**. K147-K272 and K147-K33 cross-links mapped on a c-state structure (PDB: 2C3E) and m-state structure (PDB: 6GCI). K33-K147 cross-link is mapped with Euclidean distance as SASD calculated values exceed 99 Å. **D**. Heatmap of log_2_ ratios in c2c12 isolated mitochondria treated with m-state inhibitor (BKA) or c-state inhibitor (CATR).

## CONCLUSION

Here, we have demonstrated the utility of a newly developed method for quantitative assessment of protein interactomes in cells using isobaric quantitative Protein Interaction Reporter Application of iqPIR technologies to compare interactomes of three commonly used human cell lines. These efforts demonstrated surprisingly high reproducibility and robustness of quantitative values that enable statistical filtering to reveal significant changes among HEK293, MCF-7, and HeLa cellular interactomes. Combining the quantitative cross-linking data together with DIA based quantitative proteomics identified molecular changes inside cells that were driven by and concordant with measured protein abundance level changes. The agreement in keratin cross-link levels produced in cells with measured extracted keratin protein levels demonstrated that the presented interactome approach can produce an accurate assessment of changes from inside cells. More excitingly, additional presented examples illustrate that regulation of protein conformations and protein-protein interactions within cells independent of protein abundance level changes can be uniquely detected in this way. We have identified large scale remodeling of RNA binding proteins between three cell lines that is driven in part by changing protein levels and in part by change in protein-protein interactions and confirmations. Although additional experimentation is required to better interpret these changes, conformational change of HNRPC could be either required for or a consequence of increased interaction with ROA1 that was concurrently increase in HEK293 cells. Given the roles of both HNRPC and ROA1 in mRNA binding and possibly, spliceosome function, these and other hnRNP interactome changes may be indicative of increased demand for mRNA maturation and shuttling that would be expected based on increased cell growth rates and required protein production. Indeed, our previous efforts to map interactome changes during cell growth with a mitotic inhibitor^20^ revealed an inhibitor concentration-dependent decrease in HNRPC-ROA1 K42- K350 and K147-K350 cross-link levels. Similar to our previous discovery that cross-link levels within enzyme complexes can serve as an indicator of in vivo enzymatic activity ^14^ the present observation of hnRPN interactome changes may provide novel insight on mRNA processing activity within cells.

Chromatin remodeling, glycolytic metabolism and subsequent damage by its byproducts and ADP/ATP translocation between mitochondria and cytoplasm have been extensively studied in the context of cancer^31,39,40,44–47^. Most studies have focused on changes in expression as measured by mRNA or protein levels. Quantitative cross-linking enables additional insight on these processes. Detection of changes in the chromatin remodeling complex ISWI included increased cross-link levels at a site proximal to DNA in HEK293 cells compared to HeLa, consistent with altered nucleosome interactions. Glycolysis is increased in many cancers and creates byproducts that, if not eliminated by the glyoxalase system, lead to cellular damage and eventually apoptosis^48^. The MCF- 7/HeLa interactome comparison uncovered LGUL conformational alterations near the substrate binding site and close to cysteine with inhibitory glutathionylation, independent of any change in protein levels. Finally, differences in cross-link levels in mitochondrial ADP/ATP translocase 2 and 3 indicate conformational state enrichment in MCF- 7 cells compared to HeLa. Several cross-linked species were found to correspond to specific conformational states by measuring changes in cross-link levels with state-specific inhibitors in isolated mitochondria. In general, quantitative interactome and proteome data can help identify changes in protein, conformational and protein interaction levels that shape cellular functional landscapes which can better inform cell type selection for any biological or biomedical studies.

## METHODS

### Cell culture and harvesting

HeLa cells (ATCC CCL-2), HEK293 cells (ATCC CRL-1573), and breast cancer cell line MCF-7 (ATCC HTB-22) were seeded at a density of 5 × 10^6^ cells into 150 mm culture dishes with 20 mL of Dulbecco’s modified Eagle medium (DMEM), supplemented with 10% (HeLa and HEK) and 20% (MCF-7) dialyzed fetal bovine serum (Valley Biomedical), and 1% penicillin/streptomycin (Fisher Scientific). The media was aspired and the cells were washed once with 5 mL of phosphate buffered saline (PBS). Cells were harvested with 5 mL of PBS with 5 mM EDTA for 3 min, at 37 °C. Cells were washed once with a PBS solution containing 1 mM CaCl_2_ and 1 mM MgCl_2_, and twice with PBS, pelleted each time by centrifugation at 300 × *g* for 3 min.

### Fluorescence microscopy

Fluorescence microscopy was performed as previously described, with modifications^49^. Briefly, MCF-7, HEK293, and HeLa cells were grown on Matek 2.5 cm plates with 1.5 glass coverslips until 80% confluent. Cells were washed two times with PBS, and then fixed with 1 mL of 10% formalin for 20 min at room temperature. Fixed cells were washed three times with PBS, and then blocked for 1 h in 1 mL of PBS with 5% BSA and 0.3% Triton X-100. Cells were then incubated with primary antibodies (1:250) anti-keratin 8 and (1:800) anti-keratin 18, overnight at 4 °C. Cells were washed three times with PBS before incubation with the secondary antibodies, Alexa Fluor 568 goat anti-rabbit and Alexa Fluor 488 goat anti-mouse (Life Technologies) at 1:1,000 dilutions for 1 h at room temperature. Cells were then washed three times with PBS before imaging using Cytation 5 (Biotek). Image analysis was performed using GIMP 2.10.30.

### *In vivo* cross-linking

Chemical cross-linking was performed with two independent biological replicates for comparison of HeLa and MCF- 7 cells lines, and four independent biological replicates for the comparison of HEK293 and HeLa. Cross-linking was done by individually adding 5 mM of each iqPIR cross-linker (stump heavy or reporter heavy^10^) in 500 μL of cross- linker buffer (170 mM Na_2_HPO_4_, pH 8.0) to each one of the cell suspensions. Cross-linking reactions were performed at room temperature for 30 min, with constant shaking, as previously described^10,11^. The iqPIR labeling of cells was designed as follows: (i) HeLa stump heavy *versus* HEK reporter heavy (referred here as forward), (ii) HeLa reporter heavy *versus* HEK stump heavy (referred here as reverse), (iii) MCF-7 stump heavy *versus* HeLa reporter heavy, and (iv) MCF-7 reporter heavy *versus* HeLa stump heavy.

### Sample preparation

Cross-linked cell pellets were suspended in 8 M urea solution in 0.1M NH_4_HCO_3_ buffer, pH 8.0, and sonicated 5 times using a GE-130 ultrasonic processor, with amplitude of 40% for 5 seconds. After Bradford assay for protein quantitation, 4 mg of protein from reporter heavy and stump heavy labeled samples were mixed in one 15 mL Eppendorf tube (8 mg total). This was followed by reduction with 5 mM *tris*(2-carboxyethyl)phosphine (TCEP) for 30 min, followed by a 30 min incubation with 10 mM iodoacetamide (IAA), and tryptic digestion. Peptide samples were acidified to pH 2 with trifluoroacetic acid (TFA), followed by desalting by solid phase extraction using C18 Sep-Pak cartridges (Waters) and concentration by vacuum centrifugation. Resulting peptides were suspended in 500 μL of 7 mM KH_2_PO_4_ in 30% acetonitrile, pH 2.8 and fractionated by strong cation exchange (SCX) chromatography using an Agilent 1200 series high-performance liquid chromatography system equipped with a 250 × 10.0 mm column packed with Luna 5 mm diameter, 100 Å pore size particles (Phenomenex). Peptide were separated using a binary mobile phase solvent system consisting of solvent A and solvent B (7 mM KH_2_PO_4_, 350 mM KCl, 30% acetonitrile pH 2.8) at a flow rate of 1.5 mL/min using the following gradient: 0-7.5 min 100% A, 7.5-47.5 min 95% A/5% B to 40% A/ 60% B, 47.5- 67.5 min 40% A/ 60% B to 100% B, 67.5-77.5 min 100% B, 77.5-97.5 min 100% A. A total of 14 fractions were collected and pooled as follows: fractions 6-7, 8, 9, 10, and 11-14. Cross-linked peptides from SCX fractions were further enriched using UltraLink monomeric avidin (Thermo Fisher Scientific). The enriched cross-linked peptide sample was concentrated by vacuum centrifugation and stored at -80 °C.

### LC-MS/MS analysis of cross-linked peptides

Pooled peptides from SCX fractions (6-7, 8, 9, 10, and 11-14) were resuspended in 30 μL of 0.1% (vol/vol) formic acid. Fractions were centrifuged at 16,000 × *g* for 10 min at room temperature and the supernatant of each sample transferred to an LC autosampler vial (Thermo). For the Mango LC-MS/MS analysis, 3 µL of each sample (approximately 1 µg) from each pooled fraction was injected into the nano-LC system (Thermo), coupled to a Q-Exactive Plus mass spectrometer (Thermo). Each fraction was analyzed either two or three times for technical replicates of the experiment. Each peptide fraction was then loaded onto a trap column [3 cm × 100-μm i.d., stationary phase ReproSil-Pur C8 (5-μm diameter and 120-Å-pore-size particles)] with a flow rate of 2 μL/min of mobile phase: 98% (vol/vol) LC–MS solvent A [0.1% (vol/vol) FA in water] and 2% (vol/vol) LC–MS solvent B [0.1% (vol/vol) FA in acetonitrile], and chromatographically separated using an analytical column [60 cm × 75-μm i.d., stationary phase ReproSil-Pur C8 (5-μm diameter and 120-Å-pore-size particles)], applying a 240-min linear gradient: from 95% LC–MS solvent A, 5% LC–MS solvent B to 60% LC–MS solvent A, 40% LC–MS solvent B, at a flow rate of 300 nL/min. For Mango analysis of the PIR cross-linked peptides, the mass spectrometer was set to high resolution method to data-dependent acquisition (DDA) of the 5 most intense ions with charge state of +4 to +8, with a dynamic exclusion of 30 s and isolation window of 3 Da. Each MS1 scan (70,000 resolving power at 200 *m/z*, automated gain control (AGC) of 1 × 10^6^, scan range 400 to 2,000 *m/z*, dynamic exclusion of 30 s) was followed by 5 MS2 scans (70,000 resolving power at 200 *m/z*, AGC of 5 × 10^4^, normalized collision energy of 30). For ReACT^50^ LC-MS^n^ analysis, the hybrid LQT Velos FT-ICR was set with DDA of the 6 most intense ions with charge state of +4 to +8, with dynamic exclusion of 45 s and isolation window of 3 Da. Each MS1 scan (50,000 resolving power at 200 *m/z*, mass range 500 to 2,000 *m/z*) was followed by 1 MS2 acquisition for the “on-the-fly” mass relationship check – mass precursor = mass reporter + mass peptides – (12,500 resolving power at 200 *m/z*, CID activation, normalized collision energy of 25), and 4 MS3 acquisitions of the selected ions with a 20 ppm tolerance for mass error after the mass relationship check, and with 1+ and 2+ charge states (CID activation, normalized collision energy of 35). This strategy resulted in 2 technical replicates (mango and ReACT) for each biological replicate of MCF-7 versus HeLa (forward and reverse).

### Data processing and cross-linking quantitation

For mango^51^ analysis, .ms2 files were generated containing individual precursor masses of released peptides for each spectrum and .peaks file containing all relationships within a 20 ppm tolerance. Comet^52^ (version 2019.01 rev. 4) was used to search mzXML files against the human database downloaded from UniProt^53^ on January 1^st^ 201. For MCF-7/HeLa comparison, 3,234 unique protein sequences included in the database were identified as putative cross-linked proteins through a stage 1 database approach as previously described^19^. For ReACT *in silico* analysis, MS^3^ spectra containing peptide fragmentation information was searched within 10 ppm against the same database using Comet. Then, react2csv was run on search results (*pep.xml files) to map ReACT2 results to the sequences in the database. The data were filtered to an estimated maximum false discovery rate (FDR) of 1% at the non-redundant cross-linked peptide level. The cross-linking quantitation was done as describe^16^. The cross-linking results were uploaded to XLinkDB^54^, available at http://xlinkdb.gs.washington.edu/xlinkdb/MCF-7_HeLa_iqPIR.php and http://xlinkdb.gs.washington.edu/xlinkdb/HEK_HeLa_iqPIR.php. XLinkDB uses Cytoscape^55^ to generate cross- link and protein networks, Lorikeet (https://uwpr.github.io/Lorikeet/) for spectrum view, xiNET^56^ for cross-link network display, and NGL viewer^57^ for visualization and edition of protein structures and cross-links. AlphaFold^58^ structures were uploaded to NGL viewer for cross-linking mapping.

### Data analysis and statistical analysis

A 95% confidence interval was calculated for each log_2_ ratio; cross-links with log_2_ ratios in all biological replicates (MCF-7 to HeLa comparison) or with log_2_ ratios in at least three biological replicates (HEK293 to HeLa comparison) and with each log_2_ ratio having 95% confidence less or equal to 0.5 and having at least 4 ions contributing to the ratio were considered quantified with high confidence. Student’s one-sample *t*-test was performed for each log_2_ ratio based on every peptide and peptide fragment quantified and *p*-value was assigned. Bonferroni multiple testing correction was applied based on number on all confidently quantified cross-links. Confidently quantified cross-links with Bonferroni corrected *p*-value less than 0.01 and |log_2_ ratio| ≥ 0.5 were considered significantly changing between cell lines in each comparison. Correlation, volcano plots, and density plots were generated in R using ggplot2 package^59^.

### DIA proteome quantitation

MCF-7, HeLa, and HEK293 cells were grown with three independent biological replicates in the same conditions as for cross-linking experiments. After harvesting, cells were lysed in 8M urea. Proteins were reduced with 5 mM TCEP, alkylated with 10 mM IAA, IAA andsted overnight with trypsin. Peptides were then desalted on c18 Seppak columns (Waters), dried down and resuspended in 0.1% formic acid. 1 ug of peptides was loaded on the column for each run. Peptides were separated on a 30 cm column packed with c18 reprosil, 5 um resin. Data was acquired and processed as described^60^. Briefly, gas fractionated libraries were acquired using overlapping 4 *m/z* windows using a pooled sample. Then large window (24 *m/z*) DIA data was acquired for each individual sample. The data was processed in EncyclopeDIA^61^ and output with protein intensities was used for downstream analysis in R. Protein intensities were then log_2_ transformed and median normalized. The normalized values were used for heatmaps and boxplots. *T*-test and Benjamini-Hochberg correction for multiple testing was performed using built in R functions. Adjusted p-value of 0.05 and |log_2_ FC| ≥ 0.5 were used to determine significance.

### C2c12 mitochondrial isolation and cross-linking

C2c12 cells were grown in DMEM media supplemented with 20% FBS and 1% penicillin/streptomycin. Differentiation was induced by switching to 2% horse serum when the cells were 70% confluent. Cells were collected similarly to human cell lines described previously. Mitochondria was isolated according to previously published protocol^62^. Briefly, cells were homogenized in Dounce homogenizer in isolation buffer (10 mM Tris-MOPS, 1 mM EGTA, 200 mM sucrose, pH 7.4) with 1 µM BKA or CATR purchased form Cayman Chemical, or without wither for controls. The homogenates were spun down at 600 g to get rid of cellular debris. Then supernatants were spun down at 7,000 × *g* and mitochondrial pellets were washed once. Mitochondria were cross-linked in 150 µL of cross-linking buffer in the presence of wither either BKA or CATR with reporter heavy reagent (RH). Controls were cross-linked with stump heavy (SH) reagent. Cross-linking reaction was allowed to proceed for 30 min. Supernatants were then removed by centrifugation and mitochondrial pellets were lysed. BKA or CATR treated cross-linked mitochondria were combined with control mitochondria in 1:1 ratio based on protein amount and processed similarly to cross-linked cell samples described earlier.

## DATA AVAILABILITY

The datasets generated during this study are publicly available at XLinkDB with interaction table at http://xlinkdb.gs.washington.edu/xlinkdb/PHP_SAVED_VIEWS/anna/HEK293_HeLa_and_MCF7_HeLa_iqPIR_cell_lines_interactomics.php. The mass spectrometry cross-linking data have been deposited to the ProteomeXchange Consortium via the PRIDE^45^ partner repository with the dataset identifier PXD025923. Reviewer account details: with password xgOFEtPg. DIA proteomics data has been uploaded with identifier PXD052801 (reviewer access: with password 3jsINunWPAJ4). R markdown with downstream data analysis is available at Bruce lab GitHub.

## ACKNOWLEDGEMENTS

The authors acknowledge and thank all members of the Bruce lab for helpful comments and suggestions in the course of this work and manuscript preparation. This work was supported by the US national Institutes of Health through grants: R35GM136255, R01HL144778, and R01GM086688.

## SUPPLEMENTAL MATERIALS

**Supplementary Table 1. Complete cross-linking information for HEK293/HeLa and MCF-7/HeLa datasets.** pepA,B: amino acid sequence of cross-linked peptides; accession/Uniprot A,B: protein identifiers from UniProt; modposA,B: position of the cross-linked peptide in the protein sequence; geneA,B: gene name. Log_2_ ratios of cross-links are informed in the table for the two datasets, identified as: log2ratio_forwrad, log2ratio_reverse, log2ratio_all_ions.

**Supplementary Table 2. List of significantly changed cross-links in HEK293/HeLa, MCF-7/HeLa datasets.** Cross-link: a pair of cross-linked peptides with number of cross-linked residue after each sequence; uniprotA/B: uniport identifier for each protein in a cross-link; modposA/B: position of modified lysine in a protein sequence; geneA/B: gene names for proteins in a cross- link; log2ratio/_HEK293/_MCF-7: log_2_ ratio for each comparison; pvalue/_HEK293/_MCF-7: p-value associated with cross-link ratio; HEK_to_MCF: inferred HEK293/MCF-7 log_2_ ratio calculated as log_2_ ratio of HEK293/HeLa over log_2_ ratio of MCF-7/HeLa. Cross-links were considered significant if it was quantified in at least 3 out of 5 original biological replicates for HEK293/HeLa dataset or in 2 biological replicates for MCF-7/HeLa dataset, 95% confidence of the final ratio was less than 0.5, and Bonferroni corrected *p*-value of a one-sample Student’s *t*-test for quantified ions for each cross-link was less than 0.05.

**Supplementary Table 3. Pathway analysis output from STRING.** Gene ontology molecular function (MF) and biological process (BP) significantly changed in MCF-7/HeLa and HEK293/HeLa.

**Supplementary Table 4. Cross-linking information of hnRNPs and YBOX-1 proteins for HEK293/HeLa and MCF-7/HeLa datasets.** pepA,B: amino acid sequence of cross-linked peptides; acession/Uniprot A,B: protein identifiers from UniProt; modposA,B: position of the cross-linked peptide in the protein sequence; geneA,B: gene name. Log_2_ ratios of cross-links are informed in the table for the two datasets, identified as log2ratio_all_ions, filtered for zero missing values and 95% confidence intervals for the log_2_ ratios of less than 0.5.

---

**Supplementary Figure 1.**
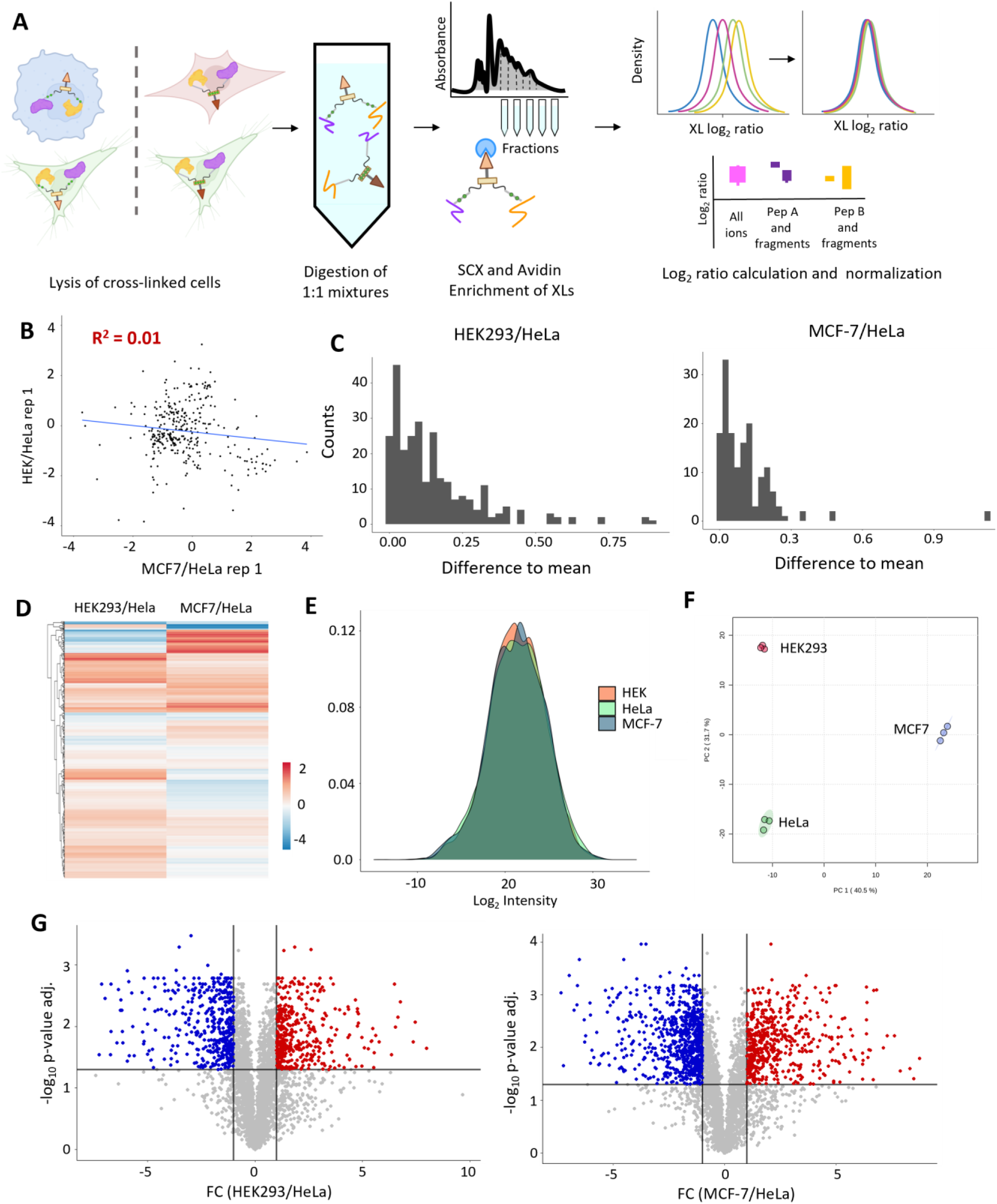
Quantitation of cross-links and proteins. **A**. Workflow for processing cross-linked cells. **B**. Correlation plot between MCF-7/HeLa and HEK293/HeLa replicates. **C**. Difference to the mean for quantitation of cross-links corresponding to same residue pairs for HEK293/HeLa (left) and MCF-7/HeLa (right) datasets. **D**. Heatmap of cross-links common between MCF-7/HeLa and HEK293/HeLa datasets. **E**. Distribution of log_2_ intensities for DIA data in all three cell lines. **F**. PCA plot based of DIA data shows clustering of biological replicates according to cell line. **G**. Volcano plots of protein level fold changes and Benjamin-Hochberg corrected *p*-values with 0.05 cutoff for significance.

**Supplementary Figure 2.**
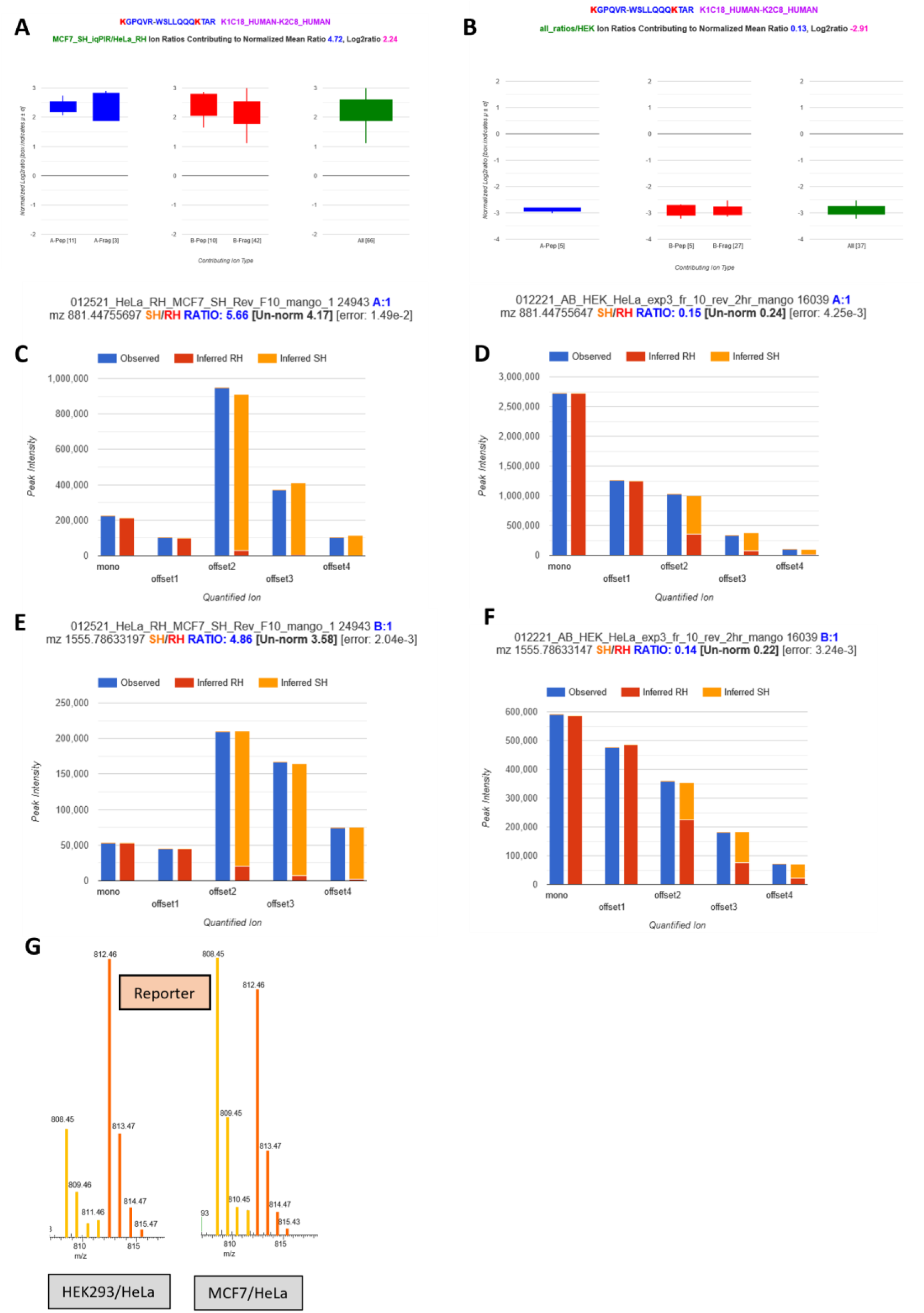
Quantitation of K1C18 K119 to K130 on K2C8 cross-link. **A, B**. Boxplots of all ions quantified in MCF-7/HeLa reverse replicate (A) or HEK293/HeLa reverse replicate (B). **C-F**. Decomposition of observed isotopic envelops of released peptides with iqPIR algorithm into each channel (RH or SH) contribution. **G**. Reporter spectra with light reporter (808) in yellow and heavy reporter (812) in orange. Ratios of the reporter channels is closer to 1 than for peptides and their fragments because of ratio compression.

**Supplementary Figure 3.**
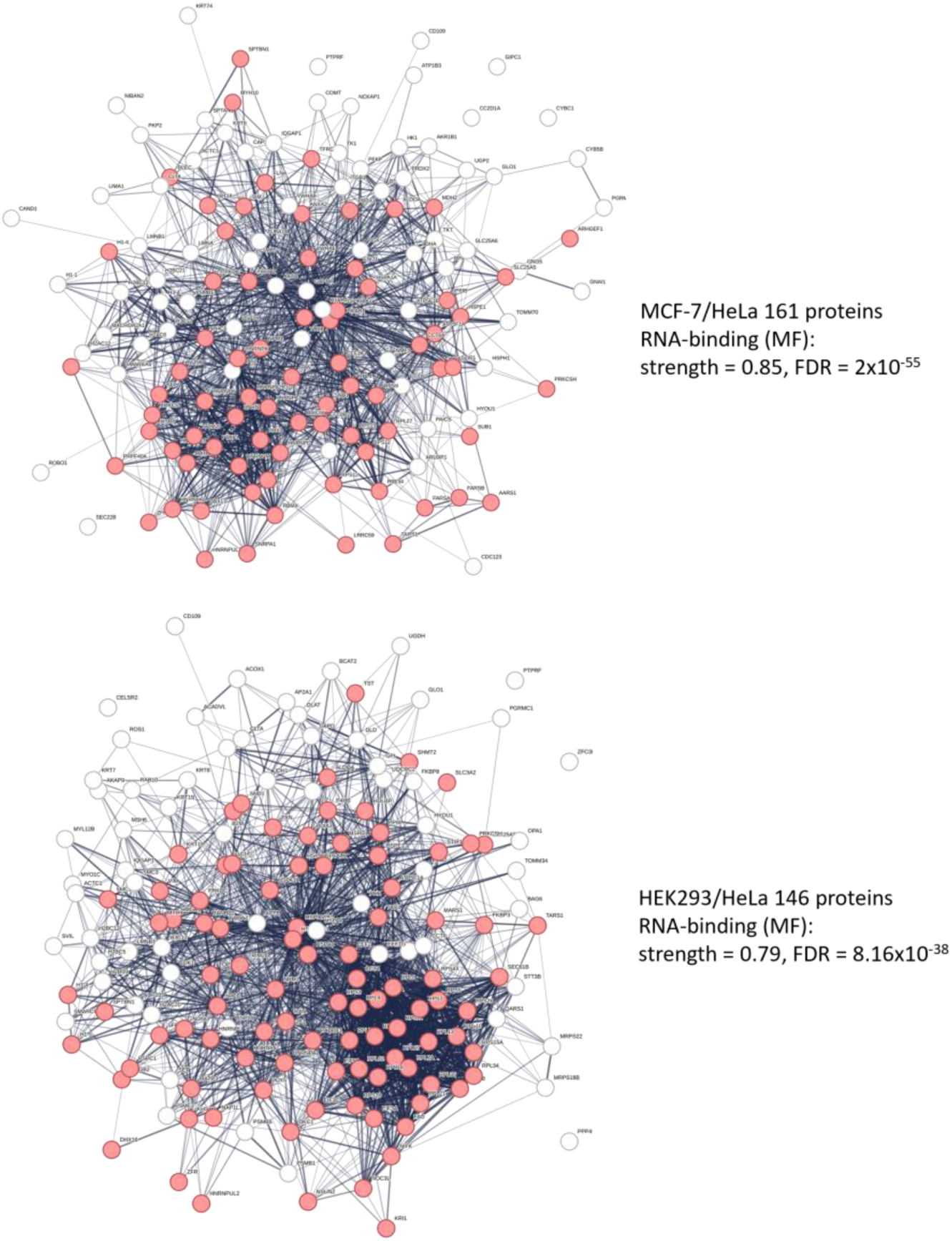
STRING network of proteins with differential cross-links. Proteins assigned to RNA binding term in molecular function (MF) annotation are red.

**Supplemental Figure 4.**
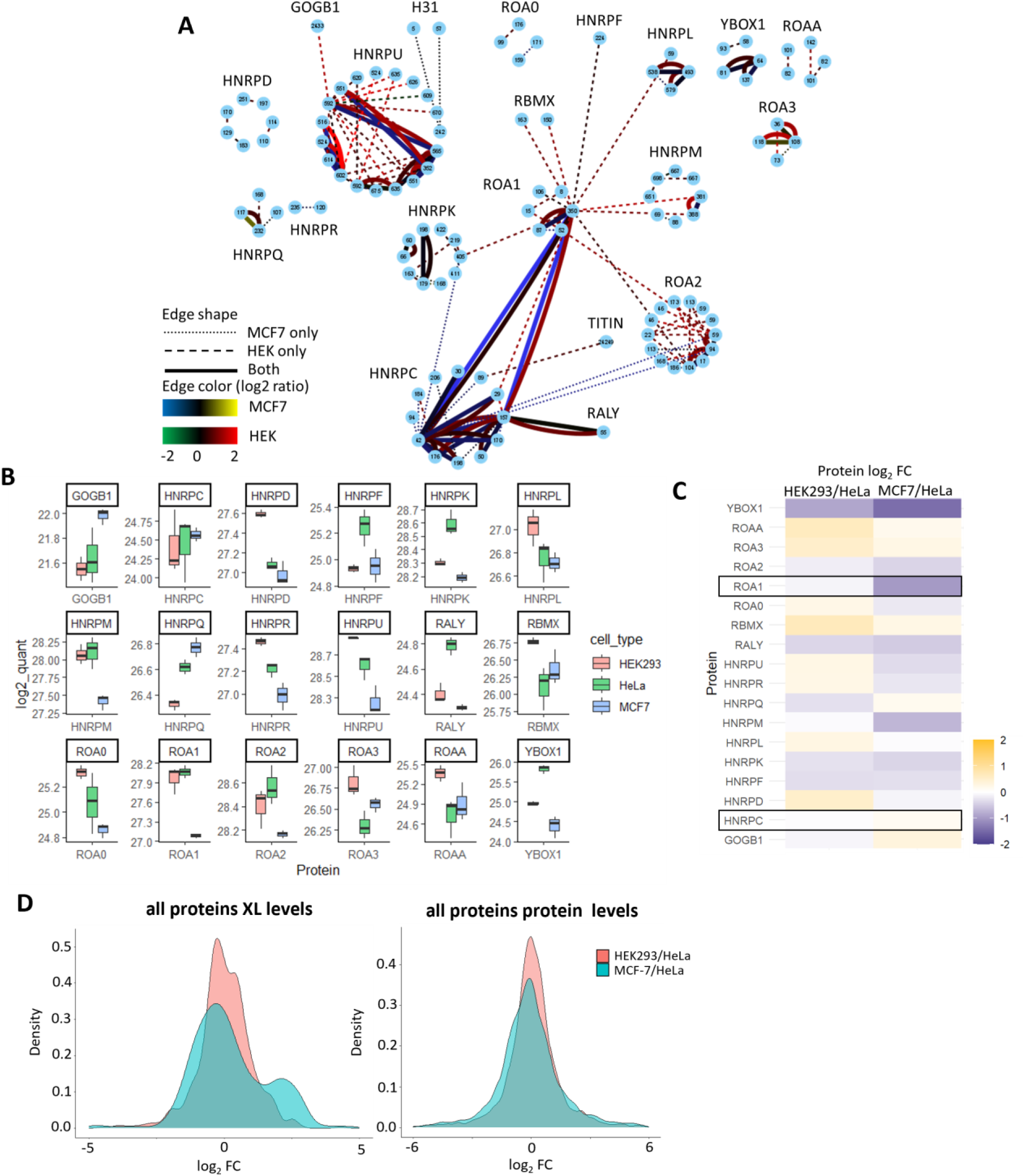
hnRNPs cross-link networks and protein quantitation. **A**. Cytoscape network for hnRNPs. Each node represents cross-linked residue with its number indicated inside the node. Edges represent detected and quantified cross- links. **B**. Boxplots comparing DIA based protein levels for hnRNPs in three cell lines. **C**. Heatmap showing log_2_ fold change in HEK293 or MCF-7 compared to HeLa of protein levels based on DIA measurements for hnRNPs. **D**. Distributions of average log_2_ fold changes for all cross-links (left) and proteins (right).

**Supplemental Figure 5.**
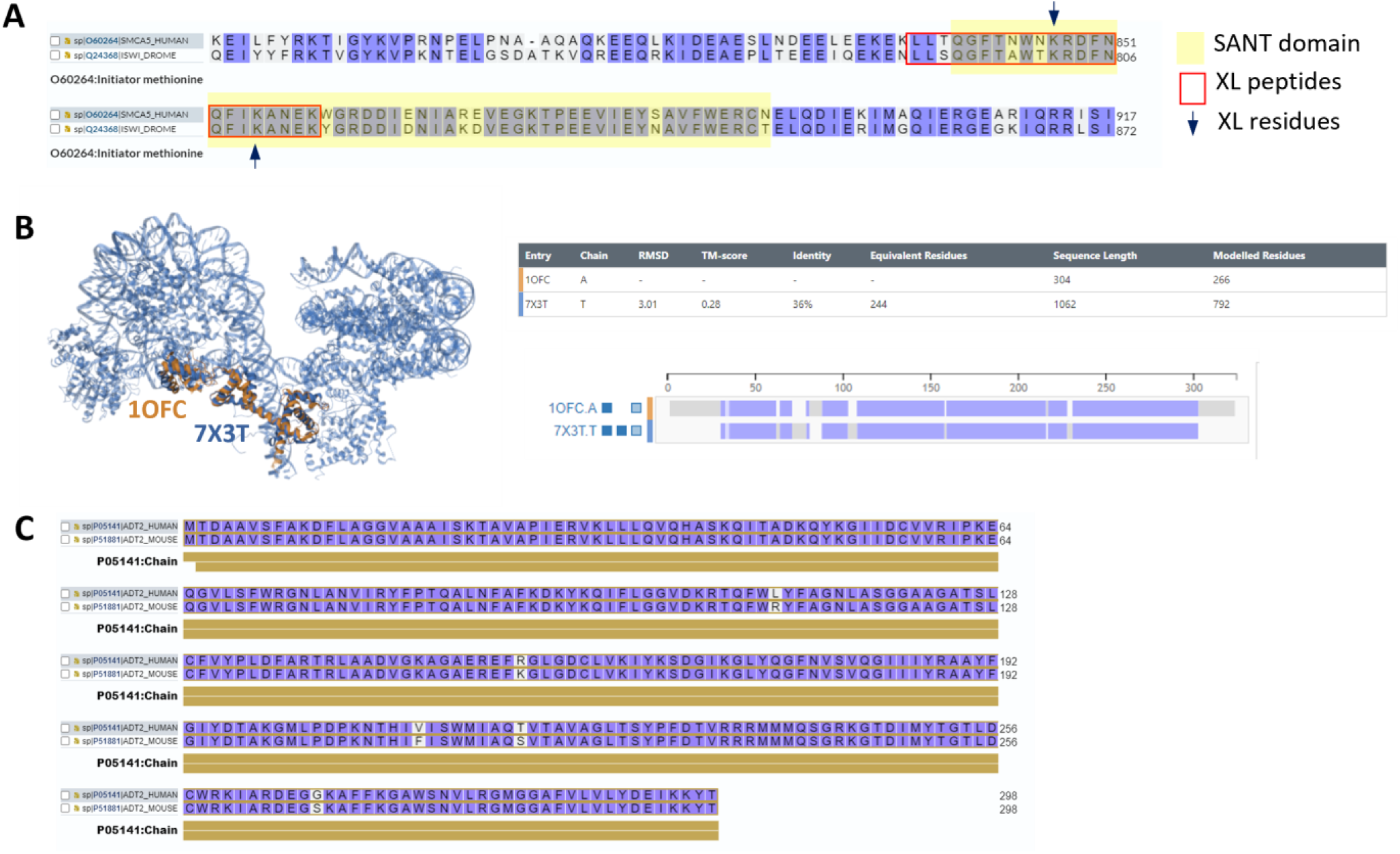
**A.** Alignment of C-terminal residues of human SMCA5 and fly ISWI. **B**. Structural alignment of fly ISWI (1OFC) and yeast ISW1a complexed with dinucleosomes (7X3T). **C**. Alignment of mouse and human sequences of ADT2 showing a high homology.

